# Optimization of regulatory DNA with active learning

**DOI:** 10.1101/2025.06.27.661924

**Authors:** Yuxin Shen, Grzegorz Kudla, Diego A. Oyarzún

**Affiliations:** School of Biological Sciences, University of Edinburgh, UK; Institute for Genetics and Cancer, University of Edinburgh, UK; School of Informatics, University of Edinburgh, UK

## Abstract

Many biotechnology applications rely on microbial strains engineered to express heterologous proteins at maximal yield. A common strategy for improving protein output is to design expression systems with optimized regulatory DNA elements. Recent advances in high-throughput experimentation have enabled the use of machine learning predictors in tandem with sequence optimizers to find regulatory sequences with improved phenotypes. Yet the narrow coverage of training data, limited model generalization, and highly nonconvex nature of genotype-phenotype landscapes can limit the use of traditional sequence optimization algorithms. Here, we explore the use of active learning as a strategy to improve expression levels through iterative rounds of measurements, model training, and sequence sampling-and-selection. We explore convergence and performance of the active learning loop using synthetic data and an experimentally characterized genotype-phenotype landscape of yeast promoter sequences. Our results show that active learning can outperform one-shot optimization approaches in complex landscapes with a high degree of epistasis. We demonstrate the ability of active learning to effectively optimize sequences using datasets from different experimental conditions, with potential for leveraging data across laboratories, strains or growth conditions. Our findings highlight active learning as an effective framework for DNA sequence design, offering a powerful strategy for phenotype optimization in biotechnology.

## 1 Introduction

The design of DNA sequences to achieve a desired phenotype is a key challenge in biotechnology. Regulatory elements, in particular, are often employed to control protein expression across many use cases, including industrial strain design [1, 2], gene therapy [3], and mRNA therapeutics [4]. Designing regulatory sequences that maximize expression often requires many iterations and domain knowledge to discover mutations that improve the desired phenotype. Moreover, the high dimensionality of genotype space together with higher order interactions between mutations can produce complex and highly nonconvex genotype–phenotype landscapes. Such landscapes are challenging to navigate experimentally and computational models are increasingly being adopted for *in silico* exploration of the genotype space [5].

Advances in massively parallel reporter assays are generating extensive sequencing and phenotypic data [6, 7, 8, 9], which enables the use of machine learning algorithms to model protein expression landscapes [10]. Such sequence-to-expression (STE) models have been developed for various regulatory elements, including ribosome binding sites [11], promoters [12, 13], transcriptional enhancers [14, 15], and 5’ untranslated regions [16, 17]. A common approach to sequence design is one-shot optimization, whereby STE models are first trained on sequencing and screening data, and then looped into a sequence optimization algorithm (Figure 1A) [18, 19].

**Figure 1:**
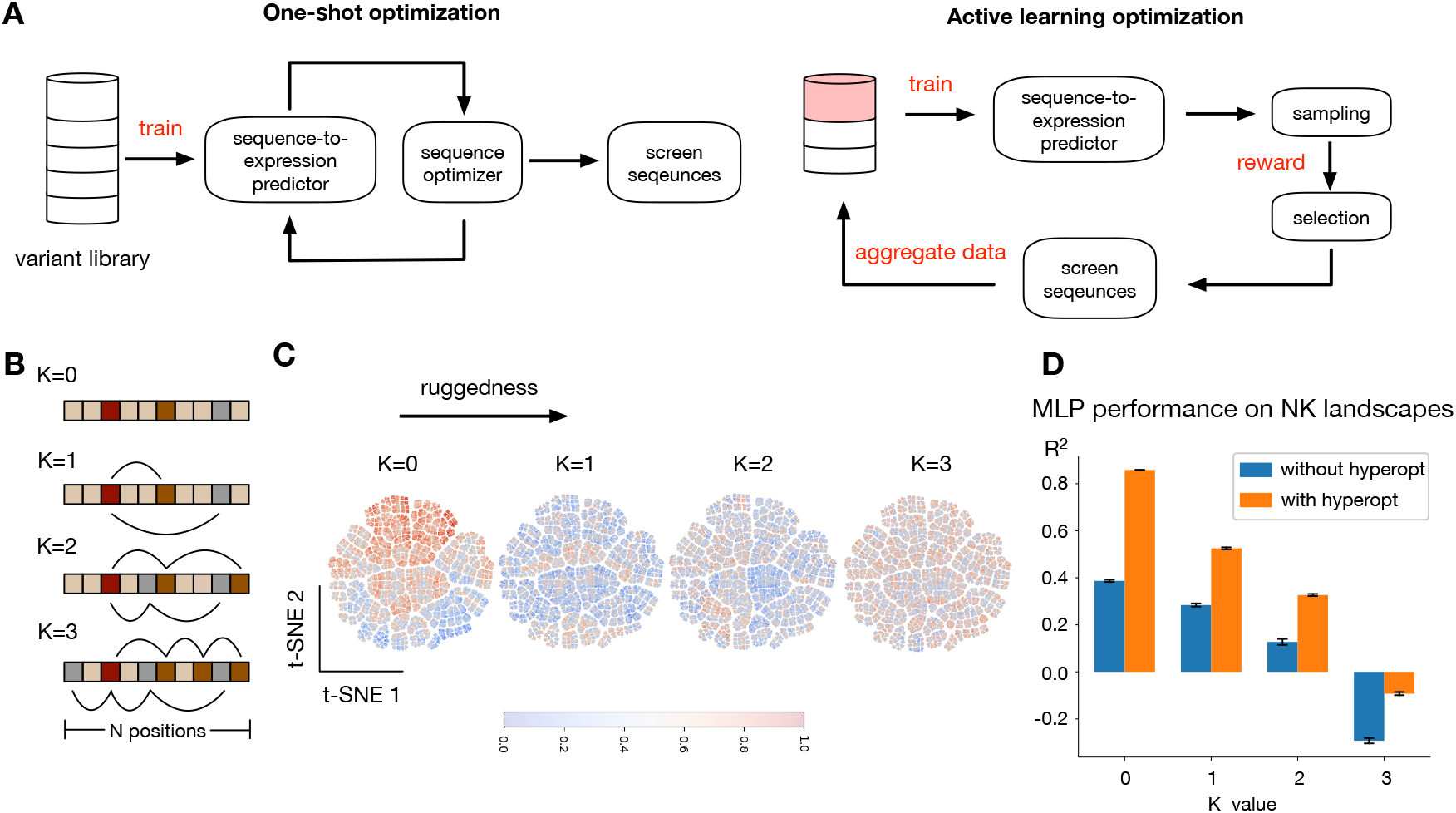
Active learning and modelling of a synthetic genotype-phenotype landscape. (**A**) Schematic of traditional one-shot sequence optimization and active learning. In active learning, experimental screening of optimized sequences is built into the optimization loop and employed to iteratively augment the training data. (**B**) NK model for fitness landscapes; *N* represents the DNA sequence length and *K* defines the number of positions that have epistatic interactions with a given position in the sequence. The value of *K* controls the ruggedness of the expression landscape. (**C**) Two dimensional t-SNE projections of the full 10nt DNA sequence space represented via one-hot encodings (1,048,576 sequences), labeled by their NK-predicted expression; t-SNE parameters no. neighbors=15 and min. distance=0.1. (**D**) Regression performance of sequence-to-expression MLP models trained on the NK synthetic data, before and after hyperparameter optimization of hidden layer size with grid search. Plots show the coefficient of determination (*R*^2^) between model predictions and simulated ground truth on a held out test set (2,000 sequences); the predictor is a feedforward neural network (MLP) trained on 2,000 sequences obtained via Latin hypercube sampling of the full sequence space. Error bars denoted the standard deviation of *R*^2^ across five random test sets of 2,000 sequences each.

The optimizer navigates the input space towards sequences with improved predicted expression, which can then screened *in vivo*. This approach has led to the discovery of improved sequences in model microbes [12, 13, 20, 21] as well as complex multicellular organisms such as drosophila, zebrafish and mice [15, 22, 23].

There are many computational techniques to traverse the STE predictions toward maxima, including gradient ascent [21], global optimization [24] and generative modelling [13, 25, 26]. However, one-shot optimization can face limitations due to the sparse and limited coverage of the sequence space employed for training—particularly as sequence length increases. In one-shot optimization, the STE model remains fixed through the sequence search, and thus optimizers can divert far away from the sequence space where the model was originally trained. This can lead to low-confidence predictions, decrease the success rate of experimental testing, and increase discovery costs. While this can be mitigated by training on larger data, the cost and complexity of acquiring large data can be a barrier in many real-world use cases [10].

Here, we explore the use of active learning for DNA sequence optimization. Active learning is a paradigm whereby models are iteratively re-trained with batches of newly acquired data (Figure 1A). This enables the adaptive selection of new sequences to measure and maximize the information gained from each round of experiments [27]. Active learning has found applications in many biological design tasks, including protein engineering [28, 29], drug discovery [30, 31], media optimization [32, 33, 34], and metabolic engineering [35, 36, 37]. We first explore the performance of active learning on a synthetic phenotype landscape produced with the classic

NK fitness model [38] adapted to nucleotide mutations. The results suggest that active learning can effectively traverse the sequence space toward increased expression even in increasingly rugged landscapes with many local optima. We then apply the methodology to an experimentally measured expression landscape containing more than 20,000,000 promoter sequences in *Saccharomyces cerevisiae*, using a highly accurate Transformer-based deep learning model as a surrogate to extrapolate expression across the whole sequence space. We demonstrate the ability of active learning to robustly find sequences with improved expression, and its ability to utilize data acquired in different growth conditions. We finally explore several performance improvements by embedding biological knowledge into the strategy for sequence sampling and selection. Our results demonstrate the utility of active learning in DNA sequence design in sparsely sampled and highly non-convex protein expression landscapes.

## 2 Results

### 2.1 Active learning on a synthetic genotype-phenotype landscape

We focus on optimization of regulatory DNA sequences using sequence-to-expression (STE) machine learning models as predictors. In a typical use case, an STE model is trained on *n* measured pairs (**x**, *y*) of genotype-phenotype associations, where **x** is a DNA sequence of length *N* and *y* is a readout of expression, typically quantified via fluorescence reporters or suitably designed screening assays that couple expression to fitness. In one-shot optimization, the STE model remains fixed and is iteratively queried by a sequence searching algorithm, typically using hill climbing or global optimization heuristics. We focus on an active learning approach (Figure 1A), whereby the STE model is re-trained during *M* learning loops, where new batches of data with *q* new samples (**x**, *y*) that are iteratively selected through the optimization loop. By careful design of a sampling-and-selection routine that balances exploration and exploitation of the sequence space, active learning can continuously improve model accuracy and navigate the predicted landscape in high-confidence regions of the STE model.

To first explore the ability of active learning to traverse highly non-convex expression landscapes, we focussed on synthetic data generated with a theoretical fitness model [39]. This allows sampling the landscape across the entire sequence space and computing global maxima as a ground truth baseline. We focussed on the NK model for fitness landscapes, because it has few parameters and produces landscapes with tuneable ruggedness. The NK model was originally developed for gene-to-gene interactions [38], and we adapted it to model epistatic interactions between positions in a nucleotide sequence (Figure 1B). In the adapted model, *N* is the number of positions in a sequence, and *K* models the number of other positions each nucleotide interacts with epistatically (i.e., the order of interaction). The parameter *K* can be tuned between *K* = 0 (no epistasis, only additive effects) and *K* = *N -* 1 to control the ruggedness, with larger *K* leading to increasingly complex landscapes with many local maxima and minima; more details on our implementation of the NK landscape can be found in the Methods.

We generated several NK landscapes for sequences of length *N* = 10, resulting in a sequence space with a total of 4^10^ = 1, 048, 576 variants. To visualize the ruggedness of the landscape, we employed the t-SNE dimensionality reduction algorithm (Figure 1C). As *K* increases, variants with more extreme phenotypes tend to appear more dispersed across sequence space, reflecting the increased ruggedness and non-convexity of the landscape. To first assess the challenge of modelling such complex landscapes, we trained feedforward neural networks on NK fitness values (Figure 1D). Models were trained on 2,000 sequences with Latin Hypercube Sampling (LHS) (Supplementary Figure S1), which represents a coverage of *∼*0.2% of the sequence space; this is substantially larger than some of the largest datasets employed in the literature. Prediction results on a held-out test set show that in the absence of epistatic interactions (*K* = 0, Figure 1D), the landscape can be regressed with reasonable accuracy. However, for *K* = 1 and above, the introduction of higher-order interactions makes the landscape significantly more challenging to regress, even after landscape-specific optimization of the neural architecture.

We built an active learning loop designed to find the global optimum of the NK fitness landscape (Figure 1A), assuming an initial set of *n* = 1, 000 variants for training that were randomly selected from the whole input space. We employed an ensemble of *r* = 10 feedforward neural networks as a machine learning regressor of the fitness landscape. At each active learning loop, sequences are sampled and selected based on the model predictions and a reward function. First, the ensemble is queried with 10, 000 sampled sequences, and the Upper Confidence Bound reward function *J*_*i*_ is calculated for each sequence *X*_*i*_:

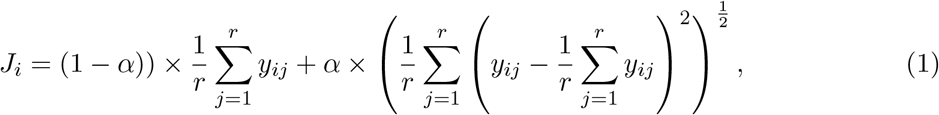

where *y*_*ij*_ is the predicted value of the the *i*th sequence with the *j*th model in the ensemble. The terms in Eq. (1) correspond to the mean and standard deviation of the predicted expression levels across an ensemble of *r* neural for each sequence. The parameter *ff* controls the balance between sequence exploration and exploration. For sequence selection, to emulate scenarios with limited data acquisition capability, we fixed the batch size to the top *q* = 100 sequences ranked by their reward function value. At each learning loop, the selected batch then goes into the evaluation step, and is added to the existing data for model re-training in the next loop.

To test the impact of the strategy employed for sequence sampling in active learning, we compared random sampling with a biologically inspired sampling based on directed evolution (DE) [40], whereby new sequences are generated by introducing mutations at *j* positions starting from sequences in the previous active learning batch. The results (Figure 2A) suggest that after four active learning loops, directed evolution sampling reaches better optima than random sampling across all levels NK ruggedness. The final-batch genotype distribution on the NK0 landscape is shown in Figure 2B, showing that active learning in tandem with directed evolution can effectively find optimal DNA sequences, in agreement with previous studies on protein fitness optimization [29].

**Figure 2:**
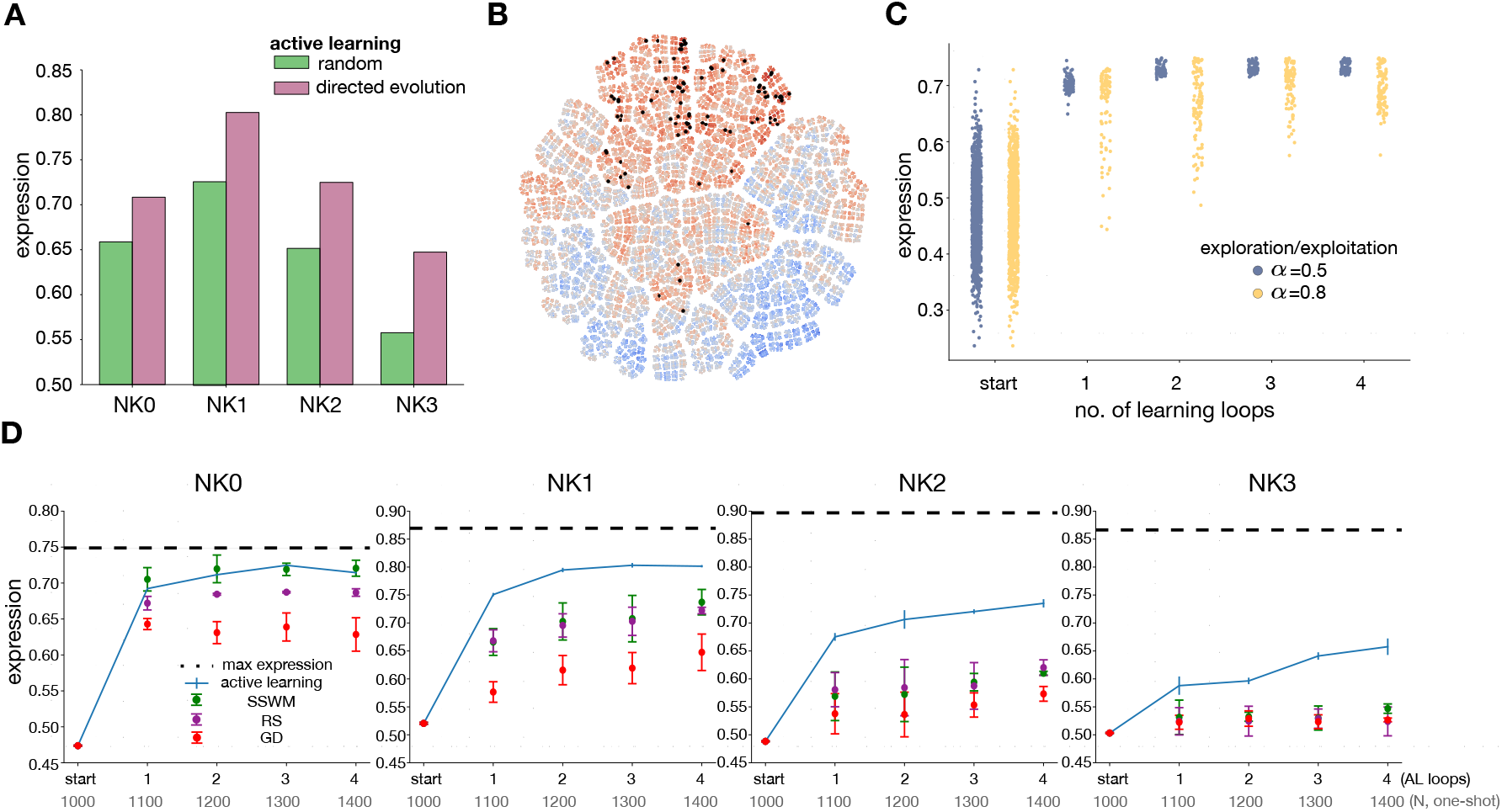
Optimization of NK genotype-phenotype landscape with active learning. (**A**) Optimized expression levels using active learning with two sampling strategies (random, directed evolution). Bars show average expression of final batch of 100 optimized sequences after four active learning loops, across NK landscapes of increasing ruggedness. (**B**) Two-dimensional t-SNE representation of the final batch of optimized sequences (black) against the ground truth expression levels (as in Figure 1C) for the NK0 landscape using directed evolution for sampling. (**C**) Expression levels for each batch of 100 optimized sequences across the active learning loops with active learning for sampling, and using different values of the exploration–exploitation parameter *ff* in the Upper Confidence Bound reward function in Eq. (1). (**D**) Comparison between and active learning and one-shot optimization in NK landscapes of increased ruggedness. Plots show average optimized expression per batch in each active learning loop using directed evolution for sampling, against several one-shot optimizers (strong-selection weak-mutation, SSWM; random sampling, RS; gradient descent, GD) ran on a multilayer perceptron (MLP) regressor. For fair comparison, the MLP was retrained on the same number of sequences as those employed for the active learning loop (*N*, one-shot). In all plots, dots and whiskers represent the mean and standard error across three replicates with resampled initial training set of 1,000 sequences, one drawn from Latin Hypercube Sampling and two from uniform sampling.

We also tested the impact of exploration and exploitation in the reward function, as their balance is crucial to ensure thorough landscape search while maintaining strong optimization performance [41]. We compared the phenotype distribution along the active learning loops in two exploration/exploitation ratio regimes using directed evolution for sequence sampling (Figure 2C). A lower ratio resulted in a higher mean phenotype distribution and with lower variance, indicating a tendency to refine predictions around already identified high expression regions while limiting broader landscape exploration.

To compare active learning with traditional optimization approaches, we implemented three one-shot optimization strategies based on random screening (RS), strong-selection weak-mutation (SSWM) and gradient descent (GD). The results (Figure 2D) across four rounds of active learning optimization suggest that active learning can outperform one-shot methods, particularly in complex landscapes with strong epistasis. We observed comparable optimization performance with SSWM in smooth landscapes (*K* = 0), but under higher-order interactions, active learning can produce better optima than one-shot optimizers. Notably, sequences selected through active learning also exhibit a clear trend of performance improvement across training loops, likely due to refinements to the STE model through re-training at each loop. We also observed large batch-to-batch improvements expression levels with active learning (Supplementary Figure S2). The success of active learning with directed evolution sampling can be attributed to its mutation from sequences that already exhibit satisfactory performance in each round of selection, which also produces STE models with improved accuracy (Supplementary Figure S3-S4).

### 2.2 Optimization of promoter sequences in yeast

To test the utility of active learning in an experimentally characterized expression landscape, we explored the optimization of promoter sequences using a large STE dataset acquired in Saccharomyces cerevisiae [12]. These data include two sets of approximately *n* = 20M and *n* = 30M promoter sequences of fixed length (*L* = 80nt), alongside expression readouts of a yfp fluorescent reporter in two different growth media. This dataset is ideal for our study because we can subsample the sequence variants to mimic different real world use cases, and test the ability of active learning to learn and transfer information across different experimental conditions (Figure 3A). To query the landscape with sequences that were not screened in the original work, we employed a transformer-based STE regressor as a surrogate for the expression landscape; this regressor was developed in the original work [12] and achieved exceptionally high accuracy on independent test sequences (Pearson *r>* 95% in both growth media).

**Figure 3:**
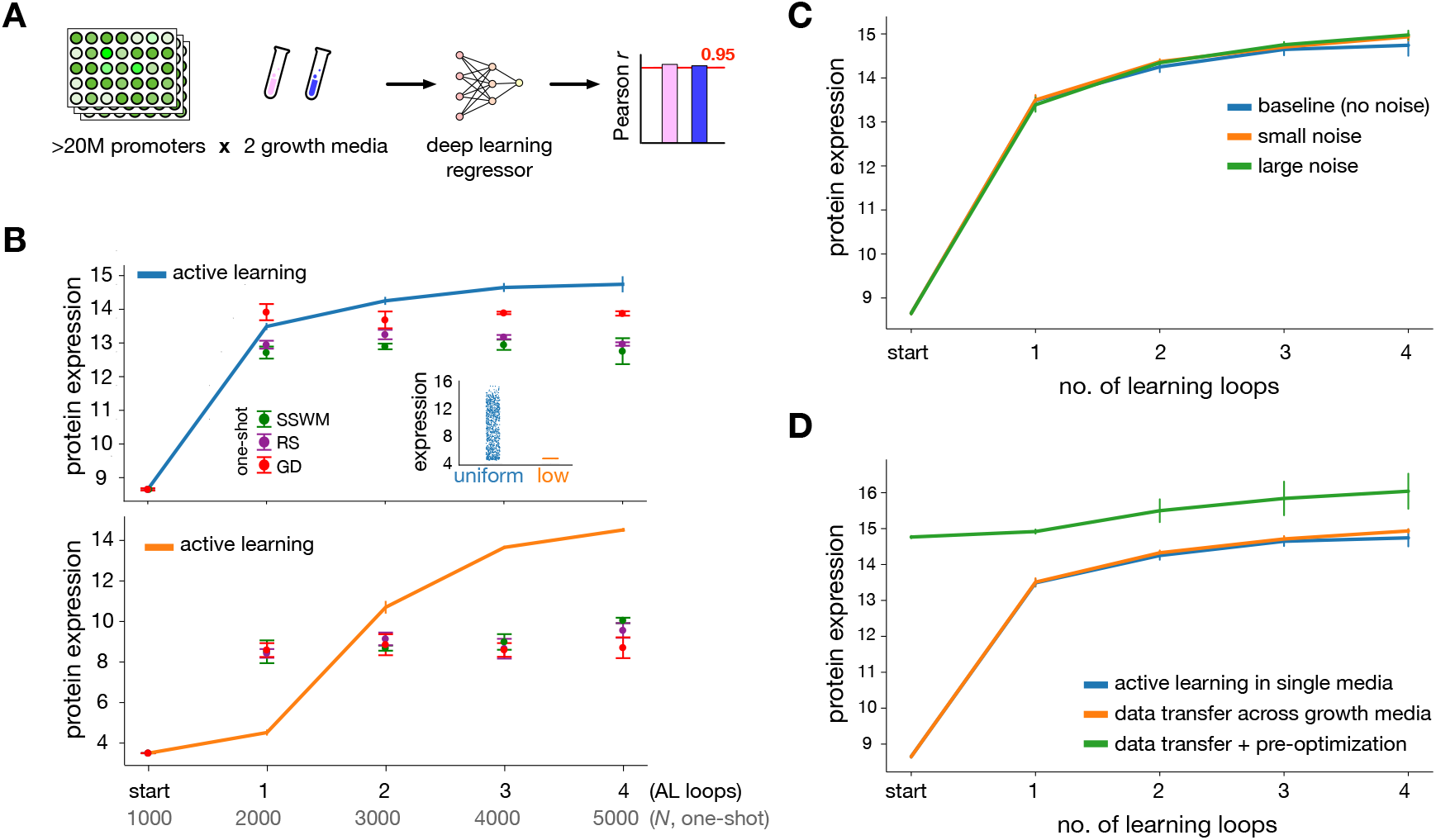
Active learning of promoter sequences in *Saccharomyces cerevisiae*. (**A**) We employed a large promoter dataset from Vaishnav *et al*, including more than 20M fully randomized promoter sequences measured in two growth media. Transformer-based models can regress the expression landscape with high accuracy (Person *r>* 0.95 in both growth media [12]). (**B**) Promoter optimization with different batches of initial data; inset shows two subsamples of *n* = 1, 000 promoters with uniform (blue) and low expression (orange); as in Figure 2D, plots show optimized expression levels across the active learning loops, compared to three one-shot optimizers on MLP regressors trained on the same number of sequences. Low expression samples were obtained by first sampling 100,000 sequences, followed by selection of sequences with lowest expression (see Methods). Dots represent the mean and whiskers show the standard error across three replicates with resampled initial training sets as in Figure 2D. (**C**) Impact of noise on active learning performance. We introduced Gaussian noise into the starting batch of measured expression levels, with zero mean and a standard deviation of 5% (orange, small noise) and 10% (green, large noise) of the ground truth value of each sequence for modelling heteroscedasticity. Dots represent the mean and whiskers show the standard error across three replicates with resampled initial training sets, as in Figure 2D. (**D**) Active learning of promoter sequences across growth conditions. We initialized the active learning loop with *n*=1,000 sequences measured in one medium, and ran the active learning optimization using iterative collection of data in a different medium. The use of sequence pre-optimization in the original medium can lead to substantial performance improvement. All active learning results employ directed evolution for sampling promoter sequences.

To mimic data scenarios encountered in applications, we considered a small initial batch of promoters in two relevant data scenarios: a case with initial promoters cover a broad range of protein expression levels, and a case in which the initial promoters are enriched for low expression (Figure 3B). This allowed testing the ability of active learning to traverse the landscape from substantially different initial starting points. We compared active learning to one-shot methods using the same total number of evaluated sequences. We employed an initial batch size *n* = 1000 promoters, with *M* = 4 active learning loops, and a batch size of *q* = 1000 top performers according to the same reward function in Eq. (1) for an ensemble of 10 feedforward neural networks evaluated at 10,000 DE-sampled sequences; we adjusted the exploration/exploitation ratio in the reward function to account for the longer sequence length compared to the previous NK landscape. In the case of uniform initial data, we observed (Figure 3B) clear batch-to-batch improvement with active learning, which outperformed one-shot methods after two active learning loops. In the case of low expression initial data, one-shot methods struggled to escape local optima at intermediate expression levels, while active learning was able to traverse expression levels close to the measured maximum. Expression distribution in Supplementary Figure S5 also shows batch-to-batch improvements in expression across the active learning loops. Despite the comparable sequence diversity in both initial batches of sequences, we observed pronounced differences in the space of sequences explored by active learning (Supplementary Figure S6).

Examination of neural network predictions against ground truth along the optimization rounds suggests that in SSWM on one-shot optimization, the fixed STE regressor tends to overshoot predictions as compared to the ground truth measurements computed with the surrogate transformer model (Supplementary Figure S7). In contrast, the STE regressor in the active learning loop progressively improves accuracy with each batch. The ability of active learning to iteratively improve model accuracy is particularly noticeable in the low expression scenario, whereby the narrow distribution of training labels leads to an initial regressor with poor generalization (Supplementary Figure S8). By iteratively supplementing the model with improved sequences, active learning can improve predictions and traverse the landscape toward better phenotypes.

To test the robustness of the active learning approach, we introduced noise to the measured expression levels for the initial batch of promoters, and thus forced the loop to start from more uncertain data. The results suggest that the active learning loop retains its ability to traverse expression towards a maximum (Figure 3C).

We sought to examine this robustness in more detail by simulating a data transfer scenario, whereby the loop is initialized with data acquired under different experimental conditions. This mimics use cases in which data from other laboratories is used to start an active learning pipeline.

We initialized the sequence optimizer with promoters measured in growth medium B, and run an active learning loop using the Transformer network pre-trained on medium A as surrogate for new measurements. The results (Figure 3D) show that the active learning loop achieves similar expression levels for both initializations. Furthermore, when initializing with sequences preoptimized in medium B, the active learning loop was able to reach a further 7.4% improvement in expression in the final batch in medium A. Overall these results suggest that active learning can effectively leverage datasets from different conditions.

### 2.3 Performance improvement with alternative sampling and selection

In this section, we explore further performance improvements via other strategies for sequence sampling and selection on the yeast promoter dataset. We first trialed alternatives to directed evolution sampling that aim to mimic other aspects of natural evolution. Specifically, we explored two alternative ways to introduce mutations in the sampled sequences (Figure 4A, see Methods for details):

**Figure 4:**
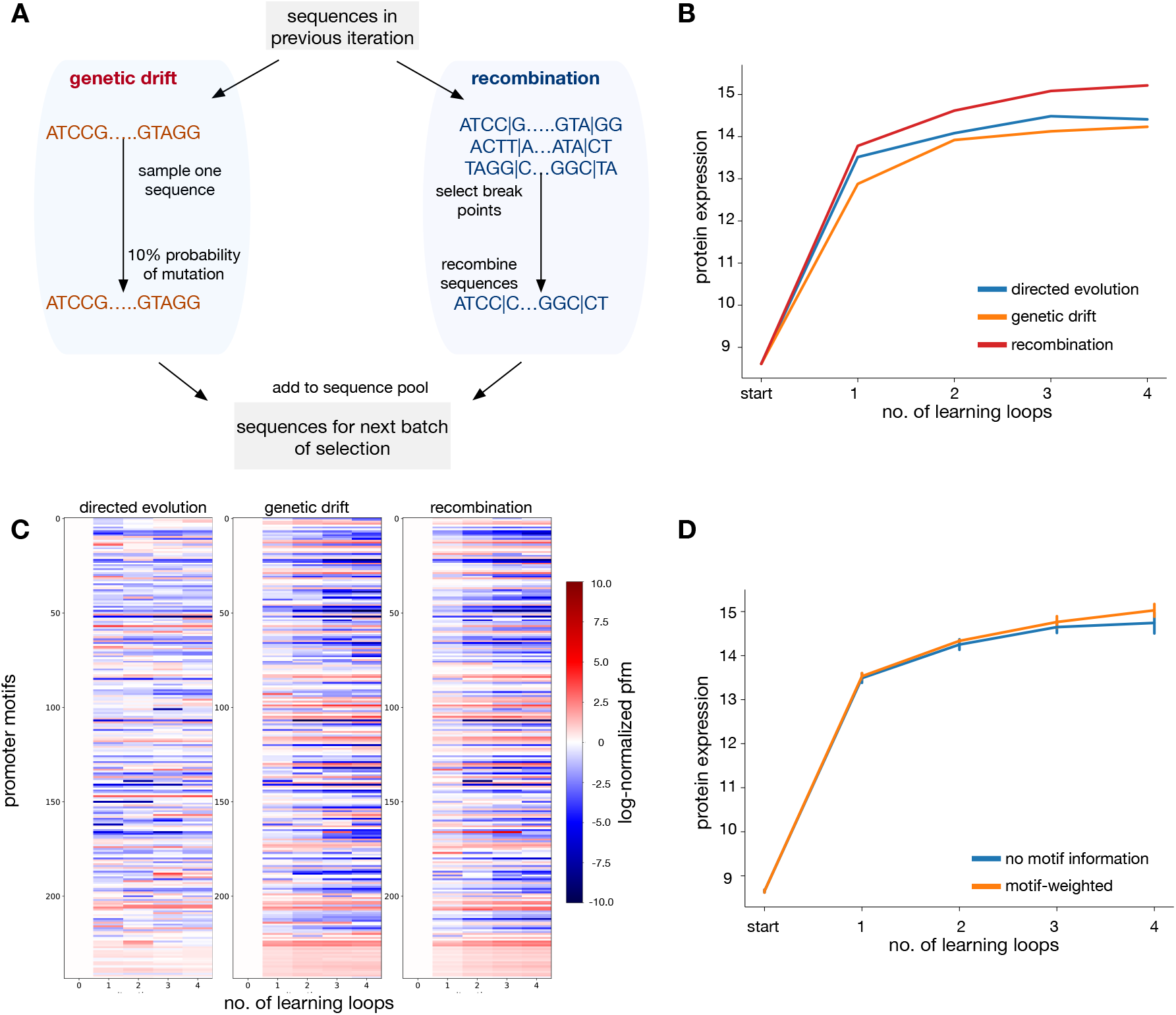
Improving active learning performance with alternative methods for sequence sampling and selection. (**A**) Two additional sequence sampling methods: genetic drift and recombination. (**B**) Comparison of active learning performance for different sequence sampling methods, initialized with a Latin Hypercube Sampling batch. (**C**) Enrichment of promoter motifs across the active learning loops for different sequence sampling strategies. Shown are average position frequency matrices (PFM) scores of 244 motifs [42] for each batch across the active learning loops; values were normalized to the PFM scores of the initial batch. (**D**) Comparison of active learning with directed evolution for sampling and selection based on a reward function with and without motif weighting, as in Eq. (2); error bars denote repeats across three initial batch of promoters.

- **Genetic drift** In this approach new query sequences are designed by assigning a probability of mutation to each site from a randomly picked sequence in the sequences from previous active learning loop. Each sequence was generated with 10% probability of mutation from a random sequence of the batch in the previous loop.
- **Recombination** We break down the sequences from the querying data of the latest learning loop, and recombine them randomly. In each loop, 10 breakpoints were randomly introduced, and the sequences from the previous loop were broken at these points and recombined.

To evaluate these sampling strategies, 10,000 sequences are sampled in each learning loop with the 3 biological sampling methods: DE, genetic drift and recombination, and the top 1000 are selected based on the reward function in Eq. (1). Figure 4B shows the performance of sampling methods across four active learning loops. The result shows the sampling methods have a substantial impact on the active learning loop. While genetic drift performs slightly worse than DE, recombination outperform DE, highlighting further potential of improving the active learning pipeline. In all three methods we observed a decreased but reasonable sequence diversity along the active learning loops (Supplementary Figure S6, Supplementary Figure S10). We also observed motif enrichment in the batches of sequences sampled by directed evolution, genetic drift and recombination in the active learning pipeline. These motifs are known to bind to specific transcription factors [42]. For each AL loop, we calculated the average motif scores across the selected batch of 1000 sequences for the 244 relevant transcription factor motifs, and compared them with the average motif score in the initial batch, which has no bias towards any motif (Figure 4C). Certain motifs were consistently enriched or depleted across all four batches and sampling methods, which suggests that active learning can capture biologically-relevant sequence features.

As another strategy to improve the active learning loop, we explored the use of introducing task-specific domain knowledge into the sequence selection step. To this end, we modified the reward function with a score that weights the presence of sequence motifs known to bind to specific transcription factors. Previous work has shown that such mechanistic information has strong correlations with expression levels [43] and helps with model generalization to new regions of the sequence space [44]. To this end, we modified the reward function by including a weight on a motif score (see (2) in Methods) mined from the YeTFaSCo database [42]. This strategy is motivated by previous work showing that most of the motifs are activators rather than repressors [43], so that maximization of the modified reward function can steer the search towards enriched with relevant motifs and hence improved expression. The results (Figure 4D) show that motif sampling improves active learning performance, indicating the effectiveness of including task-specific biological knowledge into the learning loop.

## 3 Discussion

Many applications in biotechnology and biomedicine require optimization of heterologous protein expression. One strategy is to design an expression system with highly performant regulatory DNA elements, such as promoters, enhancers or terminators, without modification to the coding sequence. Here, we demonstrated the use of active learning as a computational strategy to find regulatory sequences that improve protein expression. Using both synthetic and experimentally determined expression landscapes, our results suggest that active learning can find optimal sequences in complex expression landscapes, thanks to its ability to iteratively refine phenotypic predictions.

Active learning has been widely applied to diverse biological tasks [30, 31, 32, 33, 34, 35, 36, 37]. In the case of protein sequence design, recent studies have demonstrated that directed evolution paired with active learning can efficiently navigate protein fitness landscapes to improve enzyme function [28, 29]. Such approaches rely on iterative mutation of specific residues that are expected to impact function, such as enzymatic active sites or specific transcription factor binding motifs. In the case of regulatory DNA designed to improve expression levels, however, current experimental approaches increasingly rely on large libraries of fully randomized sequences using massively parallel reporter assays [10, 45, 46]. While this enables broader coverage of the sequence space [47], it also introduces additional challenges resulting from the ruggedness of the phenotypic landscape. Such expression fitness landscapes can have many local maxima; for example, in the case of promoter sequences such maxima may cluster around sequence regions enriched for specific transcriptional motifs.

To explore the performance of active learning on highly non-convex landscapes, we first focussed on synthetic data generated via the NK model of epistatic interactions. We show that active learning can effectively navigate such complex landscapes, which would otherwise be a substantial challenge with traditional hill climbing algorithms. We further trialed active learning on a large promoter expression dataset, using a pretrained deep learning model [12] as a surrogate for unmeasured sequences. Previous studies have explored the transfer of data across sequence-to-expression predictors models by pre-training on different experimental conditions [48], indicating that cross-condition data can provide useful information. By swapping initial data from two growth conditions, we show that active learning can robustly find optimal sequences when initialized with expression data from different experimental conditions. Notably, we found that initializing the active learning loop with pre-optimized sequences from another experimental condition led to better expression than starting from non-optimized samples from the same condition. This suggests that active learning optimization can make effective use of data acquired in different growth conditions or laboratories.

In our implementations we compared various strategies for sampling and selection of DNA sequences along the active learning loop. However, other alternatives including Bayesian optimization [49] and generative models [25, 50] could further improve performance. Moreover, in our analysis we focussed exclusively on feedforward neural networks as sequence-to-expression predictors, and further studies are needed to explore the advantages of various deep learning architectures that have shown high predictive power [51]. These and other extensions offer substantial promise for the use of active learning in DNA sequence optimization.

## 4 Methods

### 4.1 Protein expression datasets

#### Synthetic expression landscapes

To simulate synthetic genotype–phenotype landscapes with a controllable level of ruggedness, we adapted the classic NK fitness model [38] to nucleotide sequences; *N* represents the length of the DNA sequence and *K* defines the order of epistatic interactions among positions. The parameter *K* ranges from 0 to *N -* 1 and controls the ruggedness of the landscape: higher *K* values introduce higher-order epistatic interactions, increasing the ruggedness and number of local optima in the fitness landscape. We employed a previous implementation of [52] based on the code available at https://github.com/acmater/NK_Benchmarking. The generation process begins by constructing a random epistatic interaction network, such that each nucleotide position interacts with other *K* positions. Then a random fitness value is assigned to the *i*th nucleotide in the sequence based on the epistatic network. The phenotype is calculated as the average fitness values of all nucleotide positions.

#### Yeast promoter expression data

The promoter sequence ground truth models were taken from previous work by Vaishnav *et al* [12]. The original data contain yellow fluorescent protein (YFP) expression levels of 80-nt promoter sequences under two different experimental conditions: defined medium (*n* = 20M sequences) and complex medium (*n* = 30M sequences). Fluorescence measurements are in the range of [0, 18] across the whole dataset. A transformer-based predictor was trained in the original work for each experimental condition respectively, achieving high prediction accuracy (Pearson *r >* 95% in both cases). We employed these unmodified pretrained models as surrogates for experimental measurements in our active learning loops.

### 4.2 Active learning loop

#### Supervised learning of expression landscapes

We employed an ensemble of feedforward neural networks (multilayer perceptron, MLP) to regress expression levels. The ensemble enables to compute expression and estimate uncertainty of predictions for sequence sampling and selection. The ensemble consists of the top 10 performing models from a pool of 40 MLPs spanning four architectures of variable complexity and 10 random weight initializations. Model architectures are detailed in Table S1, trained with Adam optimizer and a constant learning rate of 0.001. The mean and standard deviation of predicted expressions from the 10 MLP models were used in the reward function to balance the exploration and exploitation for sequence selection.

#### Sequence sampling and selection via directed evolution

In each active learning loop, we picked a random sequence from the previous batch, and generated 10,000 randomly mutated sequences at 4 positions (NK landscape) or 10 positions (promoter landscape). The sampled sequences were then evaluated with the reward function, and the top 100 sequences on the NK landscapes or the top 1000 sequences on promoter sequence landscapes were selected for the active learning loop.

#### Reward function

To score and select sequences for the next active learning loop, we employed the Upper Confidence Bound (UCB) reward function [41] specified in Eq. (1). To determine an appropriate value for the exploration/exploitation parameter *ff*, we performed a grid search with a step size of 0.1 on the NK0 landscape (Supplementary Figure S9). The optimal value was found to be *ff* = 0.7 which was used consistently across the four NK landscapes. For the promoter datasets, we scaled *ff* by a factor based on the sequence length difference 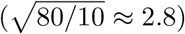.

**Implementation** Our implementation follows the following pseudocode:

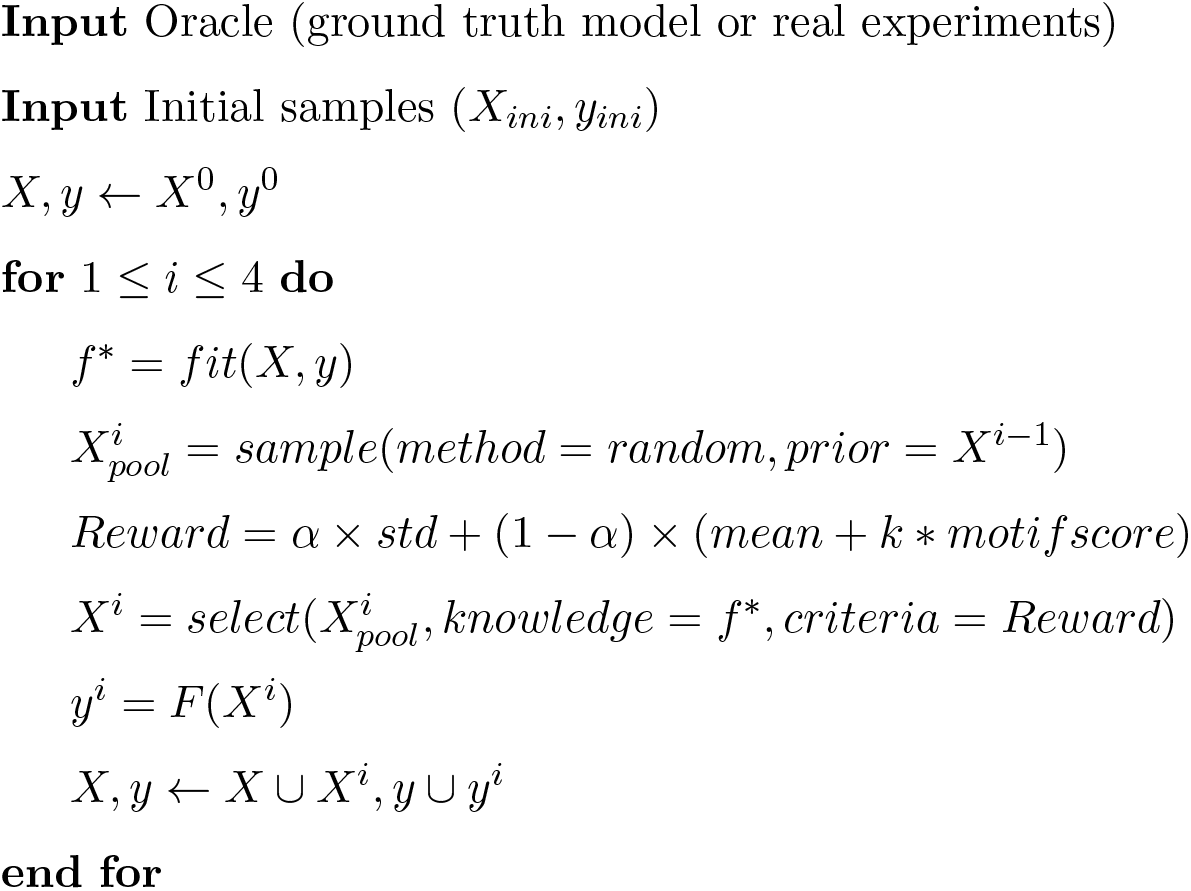

### 4.3 One-shot optimization

For fair comparisons with active learning in Figure 2–3, one-shot optimizations were computed using MLP regressors trained on the same number of sequences as in each active learning loop. The MLP architecture was selected with 5-fold cross-validation across the same architectures in Table S1, trained with Adam optimizer and a constant learning rate of 0.001. Three methods were applied consistently across the NK landscapes and the promoter sequence landscapes, detailed next.

#### Random sampling (RS)

For RS, 100,000 sequences were randomly generated and scored using the MLP model. The top 100 sequences based on predicted expression were selected.

#### Gradient descent (GD)

For GD, sequences were iterated over 200 steps to reach a higher predicted expression. At each step, MLP model gradients were estimated via finite differences and used to update the sequence in the direction that improved the model output, and the final sequence after 200 iterations was selected. The full procedure was repeated independently for 100 random starting sequences to generate 100 optimized sequences.

#### Strong-selection weak-mutation (SSWM)

For SSWM, sequences were generated following 200 generations for the NK landscapes (100 generations for the promoter landscapes) of evolution and selection to reach a higher predicted expression. In each generation, mutations were introduced at a low rate on a small population of 10 sequences, and the mutated sequences were evaluated using the MLP model. The population of next generation consisted of the top 5 sequences and 5 single-mutant variants of the current best sequence. As with GD, the full procedure was repeated for 100 times to generate 100 optimized sequences.

### 4.4 Motif enrichment and motif sampling

For the results in Figure 4D, we included additional motif score to the reward function:

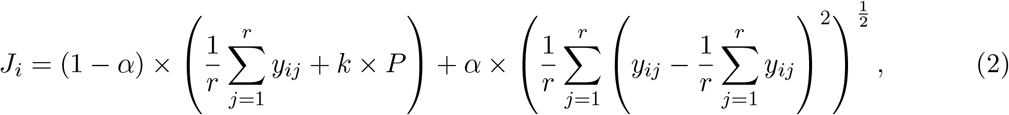

where *r* is the number of MLP predictors in the ensemble, the parameter *ff* controls the balance between sequence exploration and exploration, *P* is the sum of the motif score, and the *k* is a penalty on the motif score; we employed parameters *ff* = 0.3 and *k* = 0.66. The motif score *P* was computed from predicted binding probabilities between transcription factors (TF) and sequence motifs based on earlier work[44]. The 244 position frequency matrices (PFM) were obtained from the YeTFaSCo database [42], which provides the occurrences of each nucleotide at each position of 244 TF-specific motifs. Each PFM has dimensions of 4 × 𝓁_*i*_, where 𝓁_*i*_ is the motif length. For each promoter sequence, we compute the binding probability for each motif *P*_*i*_ by integrating over all sequence positions:

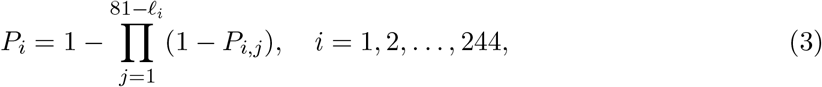

with *P*_*i,j*_ being the probability of TF binding starting from the *j*^th^ nucleotide in the 80-nucleotide sequence, derived from the corresponding PFM. The final value *P*_*i*_ is the probability that the *i*^th^ TF binds to at least one site within the entire sequence.

### 4.5 Data transfer across growth conditions

For our data transfer analysis (Figure 3D), we employed the two pre-trained transformer models (medium A, medium B) from the original data source [12] as surrogetes for ground truth data. For the data transfer across media, we first initialized the active learning loop with randomly sampled *n* =1,000 sequences measured in medium B. We then ran the active learning optimization on medium A using iterative collection of data from measurements in that medium. For data transfer with pre-optimization, we followed the same process but pre-optimized the *n* =1,000 initial sequences for medium B, and then ran the active learning loop for medium A. The pre-optimization process is a separate rounds of four active learning loops on medium B, and the final batch of 1000 sequences with highest expression in medium B was emlpoyed to initialize the loop in medium A. The benchmark (Figure 3D, blue) is the same as Figure 3B (top).

## Code availability

Python code for model training and evaluation will be made available in an open repository.

## Acknowledgments

Y.S. was supported by the UKRI Biotechnology and Biological Sciences Research Council (BB-SRC) grant number BB/T00875X/1. D.A.O. was supported by the UKRI Centre for Doctoral Training in Biomedical AI (EP/S02431X/1), and the Engineering Biology Mission Award CYBER (BB/Y007638/1).

