## Supplementary Figures for "Optimization of regulatory DNA with active learning"

### Supplementary Figures and Tables

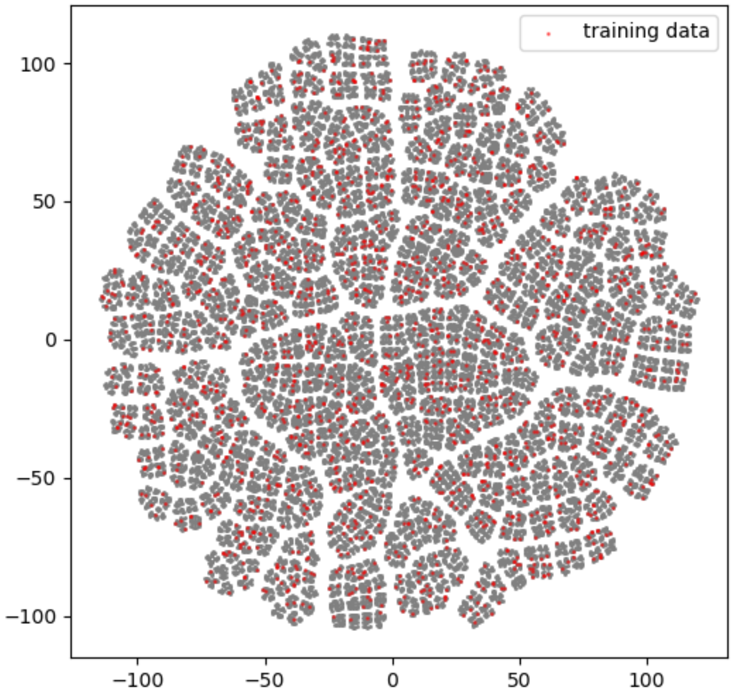


Figure S1: **Distribution of 2000 training data points for the MLP in the genotype space.** The grey points in the t-SNE plot show the whole NK sequence space of 1,048,576 sequences. 2000 sequences with LHS are used for MLP training and labeled with red in the genotype space.

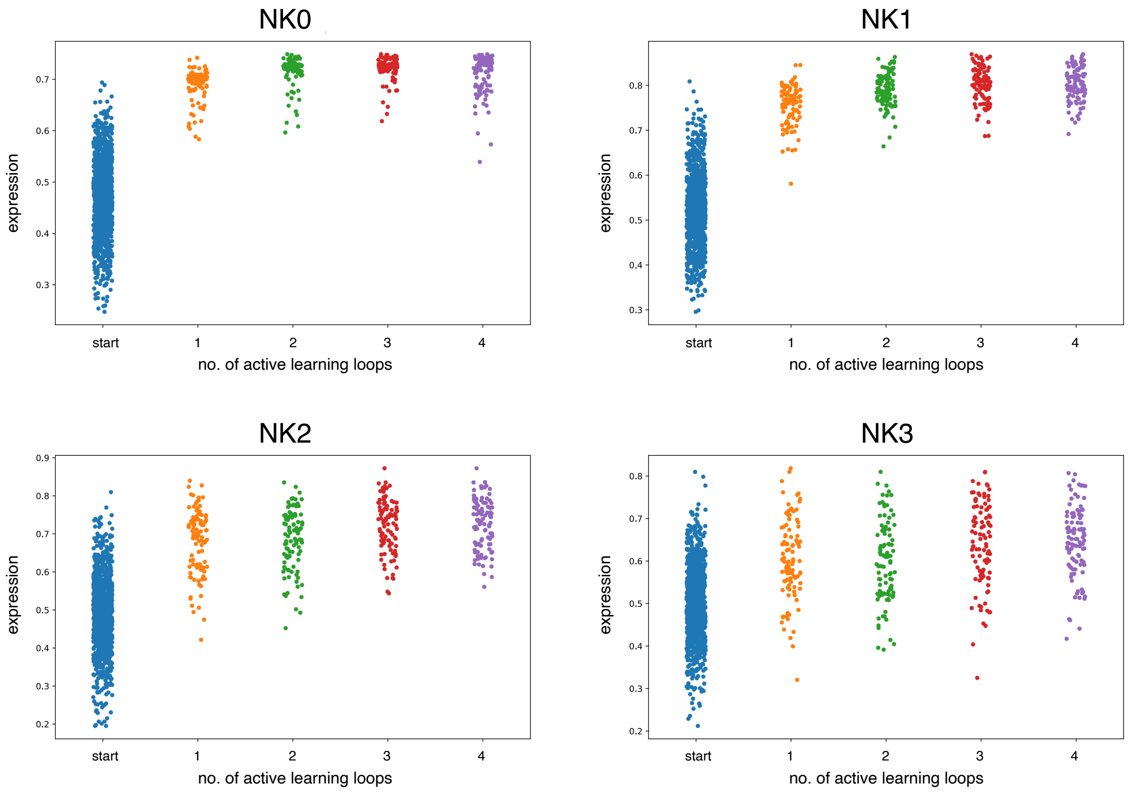


Figure S2: **Strip plot of active learning performance on NK landscapes.** The strip plots show the distribution of optimized expression levels using directed evolution as a sampling strategy for NK landscapes of increased ruggedness.

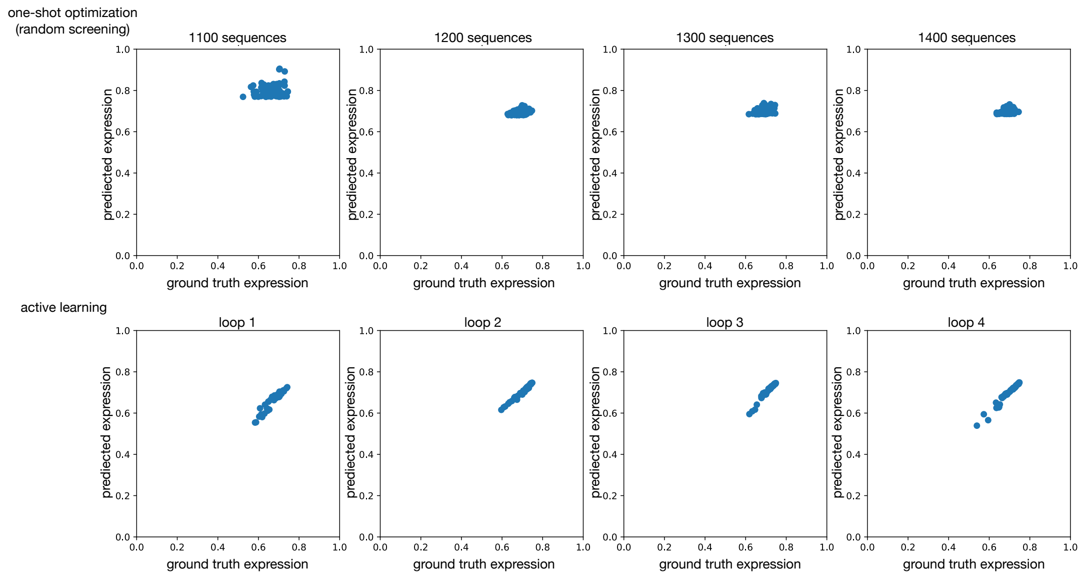


Figure S3: **Scatter plot of predicted vs ground truth expression on NK0 landscape.** The upper plot shows the scatter of random screening with one-shot optimization, and the bottom plot shows the scatter of active learning with directed evolution sampling for NK0 landscape. For the four one-shot optimization cases with different number of data or four active learning loops, the ground truth expression for each selected sequence is plotted against the predicted phenotype value by the model.

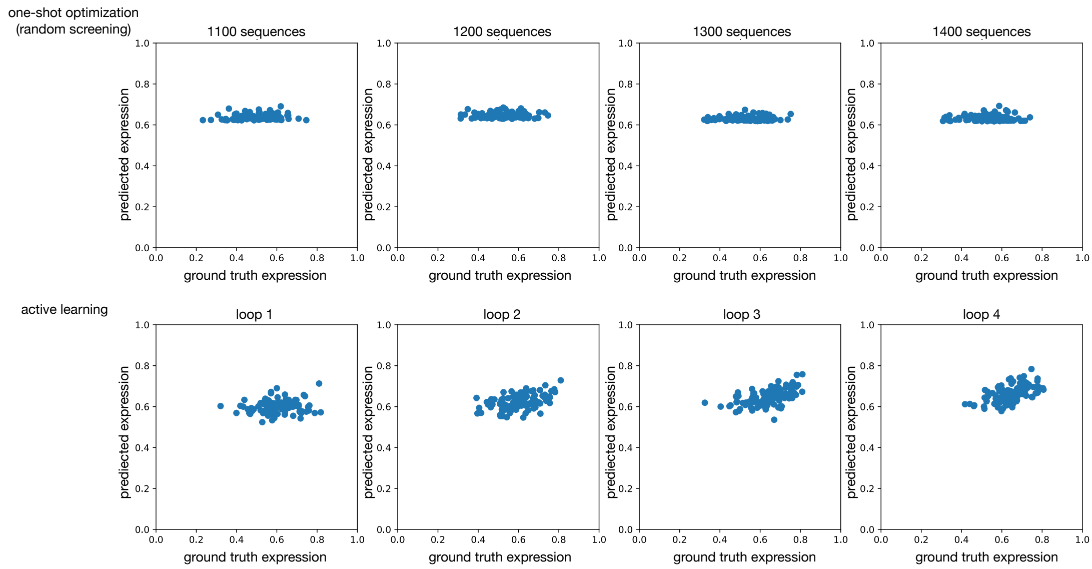


Figure S4: **Scatter plot of predicted vs ground truth expression on NK3 landscape.** The upper plot shows the scatter of random screening with one-shot optimization, and the bottom plot shows the scatter of active learning with directed evolution sampling for NK3 landscape. For the four one-shot optimization cases with different number of data or four active learning loops, the ground truth expression for each selected sequence is plotted against the predicted phenotype value by the model.

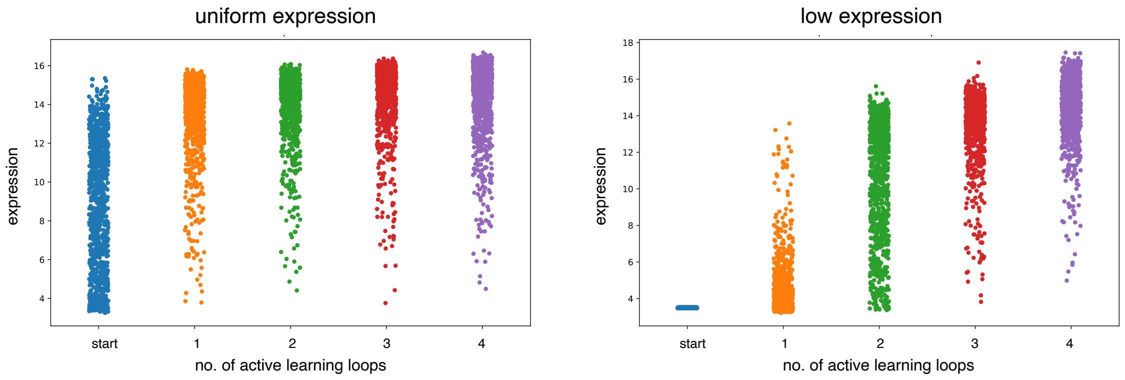


Figure S5: **Strip plot of active learning performance on promoter landscape.** The strip plots shows the distribution of optimized expression levels using directed evolution as a sampling strategy for promoter landscape. The left panel is the active learning pipeline from 1000 uniform expression promoter sequences; the right panel is the active learning pipeline from 1000 low expression initial promoter sequences.

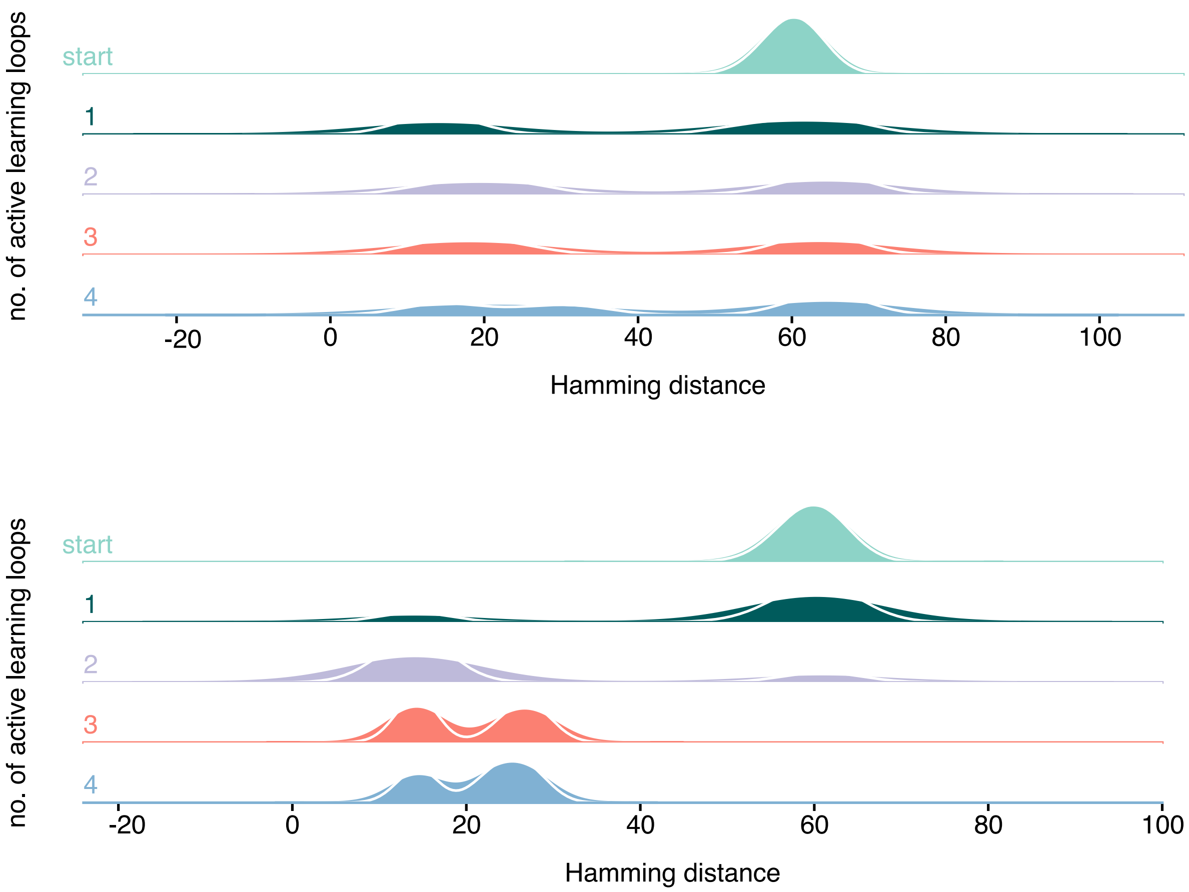


Figure S6: **Hamming distance plot in active learning pipeline for sequence optimization with directed evolution sampling.** The Hamming distance distribution between different sequences in each batch shows the sequence diversity of the optimized sequences. The upper panel shows the HD distribution of active learning loops starting with uniform initial distribution sampled by LHS, and the lower panel show the HD distribution of active learning loops starting with low expression initial sequences sampled by LHS.

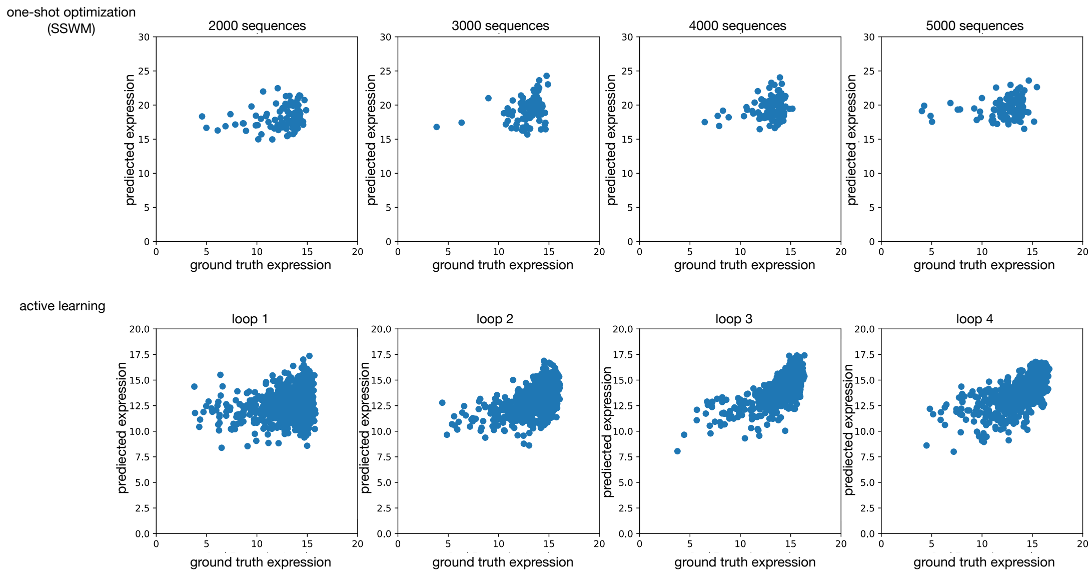


Figure S7: **Scatter plot of predicted vs ground truth expression on promoter landscape from uniform distribution.** The upper plot shows the scatter of one-shot optimization with SSWM, and the bottom plot shows the scatter of active learning with directed evolution sampling for the yeast promoter landscape from uniform expression sequences. For the four one-shot optimization cases with different number of data or four active learning loops, the ground truth expression for each selected sequence is plotted against the predicted phenotype value by the model.

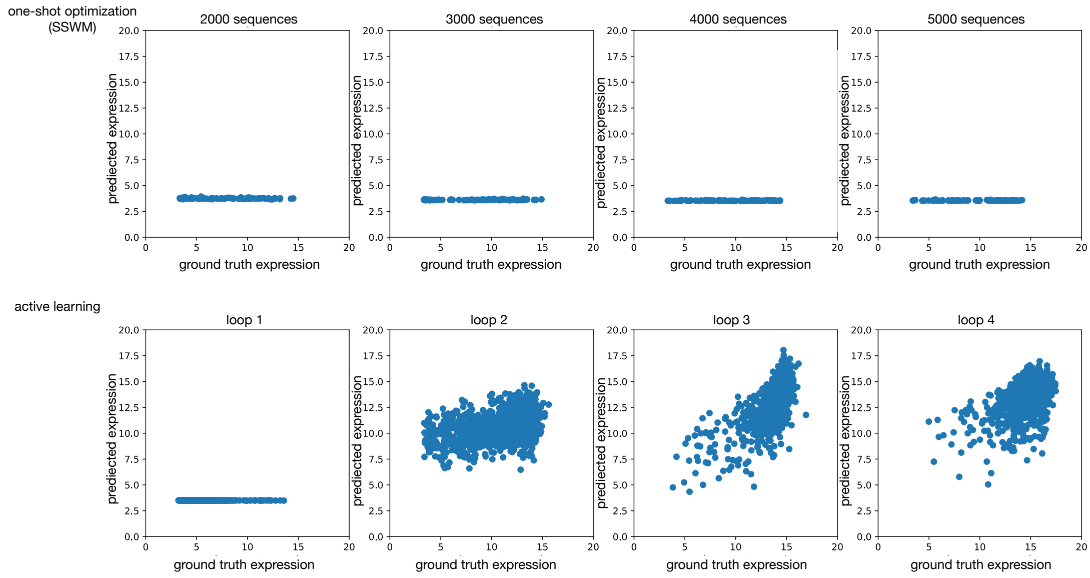


Figure S8: **Scatter plot of predicted vs ground truth expression on promoter landscape from low expression.** The upper plot shows the scatter of one-shot optimization with SSWM, and the bottom plot shows the scatter of active learning with directed evolution sampling for promoter landscape from low expression sequences. For the four one-shot optimization cases with different number of data or four active learning loops, the ground truth expression for each selected sequence is plotted against the predicted phenotype value by the model.

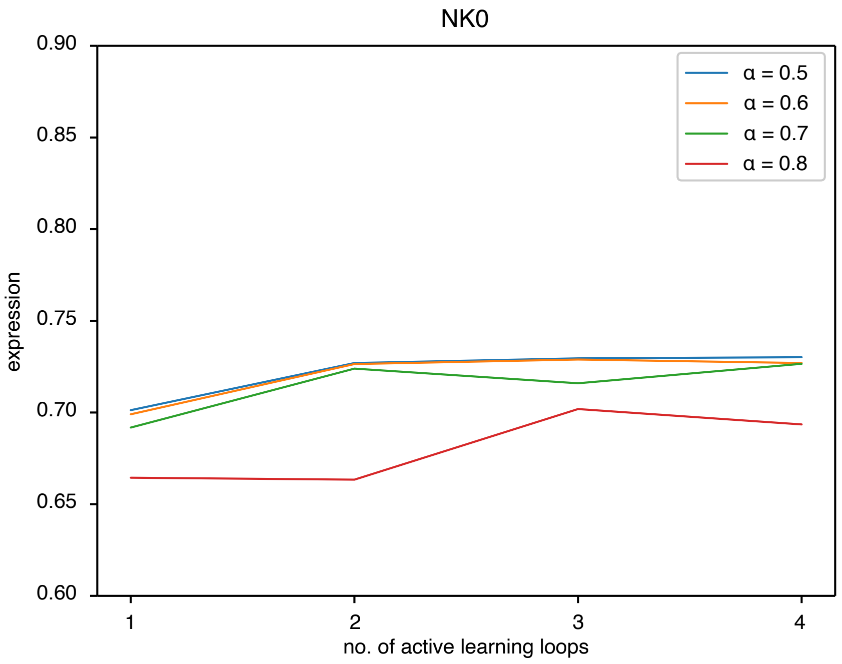


Figure S9: **Grid search of exploration factor for the NK landscape.** Performance of active learning with different exploration factor $\alpha = 0.5, 0.6, 0.7, 0.8$ in the Upper Confidence Bound reward function in Eq. (1).

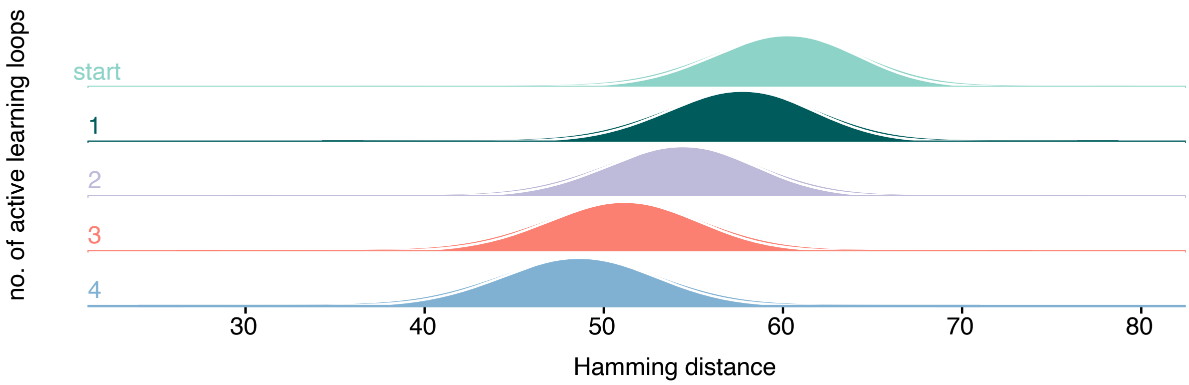


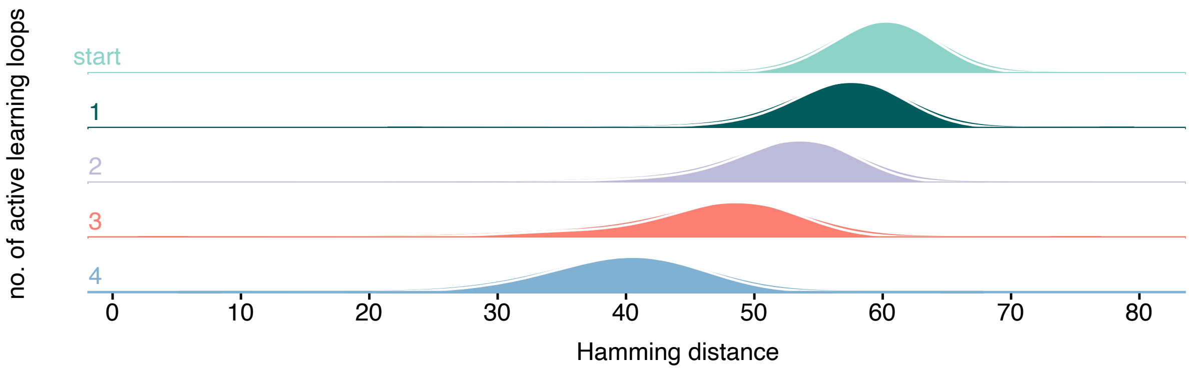


Figure S10: **Hamming distance (HD) distribution in active learning for sequence optimization with different biological sampling methods started from uniform distribution by LHS.** The Hamming distance distribution between different sequences in each batch shows the sequence diversity of the optimized sequences. The upper panel shows the HD distribution of active learning loops with genetic drift; and the lower panel show the HD distribution with recombination.

Table S1: **Hyperparameters for different MLP structures for one-shot optimization and ensemble models in active learning loop.**

| Model | no. neurons (layer 1) | no. neurons (layer 2) | no. neurons (layer 3) | no. neurons (layer 4) |
|---|---|---|---|---|
| 1 | 10 | 100 | 100 | 20 |
| 2 | 100 | 100 | 20 | / |
| 3 | 40 | 10 | / | / |
| 4 | 10 | / | / | / |
